# Covalent Flexible Peptide Docking in Rosetta

**DOI:** 10.1101/2021.05.06.441297

**Authors:** Barr Tivon, Ronen Gabizon, Bente A. Somsen, Peter J. Cossar, Christian Ottmann, Nir London

## Abstract

Electrophilic peptides that form an irreversible covalent bond with their target have great potential for binding targets that have been previously considered undruggable. However, the discovery of such peptides remains a challenge. Here, we present CovPepDock, a computational pipeline for peptide docking that incorporates covalent binding between the peptide and a receptor cysteine. We applied CovPepDock retrospectively to a dataset of 115 disulfide-bound peptides and a dataset of 54 electrophilic peptides, for which it produced a top-five scoring, near-native model, in 89% and 100% of the cases, respectively. In addition, we developed a protocol for designing electrophilic peptide binders based on known non-covalent binders or protein-protein interfaces. We identified 7,154 peptide candidates in the PDB for application of this protocol. As a proof-of-concept we validated the protocol on the non-covalent complex of 14-3-3σ and YAP1 phosphopeptide. The protocol identified seven highly potent and selective irreversible peptide binders. The predicted binding mode of one of the peptides was validated using X-ray crystallography. This case-study demonstrates the utility and impact of CovPepDock. It suggests that many new electrophilic peptide binders can be rapidly discovered, with significant potential as therapeutic molecules and chemical probes.

## Introduction

While small molecules are traditionally used for targeting specific proteins, some targets are notoriously difficult to drug using small molecules owing to the size and flatness of their surface and the lack of natural substrates. Examples of difficult protein targets include sites of protein-protein interactions, shallow allosteric pockets, transcription factors and DNA binding proteins in general^[1]^.

A common approach to address such targets is to use peptides, which cover a much larger surface area while engaging with so-called “hot-spots’’ on the target’s binding surface^[2,3]^. Such peptides can interact with their targets with high specificity and biologically relevant affinity. Although not typically compliant with Lipinski’s rule of five^[4]^, recent chemical methodologies have been shown to improve pharmacokinetic properties of peptides, such as bioavailability, permeability and *in vivo* stability using tools such as N-methylation, cyclization and incorporation of D-amino acids^[5–8]^.

However, discovering high-affinity peptide binders that overcome the inherent binding difficulties of large protein surfaces and compete with native cellular interactions^[9]^ remains a challenge. While longer peptides may be able to interact with a larger number of non-adjacent protein hot-spots increasing the binding enthalpy, this is accompanied by higher entropic cost, which often lowers the overall binding affinity. Additionally, longer peptides typically present inferior pharmacokinetic profiles.

A possible strategy to address this challenge is to introduce covalent binding between the peptide and its target, which can significantly improve the peptide potency^[10,11]^. Furthermore, covalent binders can exhibit a longer duration of action and exceptional selectivity against homologous interactors when targeting non-conserved protein nucleophiles^[12]^. There have been several reports of peptide binders that were functionalized with an electrophile to target a nucleophile at the peptide receptor binding site^[13–22]^, demonstrating the great potential of covalent peptides as drugs and chemical probes.

Despite this potential, the number of covalent peptides that have been reported is substantially lower compared to covalent small molecules. Optimization of such peptides is typically based on an iterative approach and involves only limited diversity of electrophiles and their positions. Although several automated tools have been developed to aid the modeling and design of non-covalent peptides^[23–28]^, to the best of our knowledge, no available general peptide-specific tool currently incorporates covalent binding between the peptide and the receptor.

Here, we present the computational pipeline CovPepDock, that extends Rosetta FlexPepDock^[23]^, a previous protocol for modeling non-covalent protein-peptide interactions. The CovPepDock protocol includes electrophiles as part of the peptide and models the newly formed covalent bond with a cysteine residue in the target protein. We benchmarked CovPepDock against 115 structures of disulfide-bound peptide-protein complexes and 54 structures of complexes in which an electrophilic peptide covalently binds a receptor cysteine. The protocol identified the correct binding mode with near-native accuracy in 89% and 100% of the cases, respectively.

On the basis of CovPepDock, we developed a general protocol to design covalent peptide binders that target peptide-protein or protein-protein interfaces. We chose to test the performance of the protocol against the 14-3-3σ/Yap1 protein complex. The seven 14-3-3 isoforms play crucial roles in regulating various cellular processes such as cell division, gene expression and apoptosis^[29]^. They perform their function by binding to phosphorylated target proteins such as transcription factors and kinases, to modulate various aspects of their function such as cellular localization, enzymatic activity and DNA binding affinity^[30]^. The 14-3-3σ isoform harbors a non-conserved cysteine at position 38, close to the phosphopeptide binding site (Table S1). Since this cysteine is unique in the 14-3-3 family, it can be used for selective targeting of 14-3-3σ interactions using electrophile-containing peptides or peptidomimetics. Among the binding partners of 14-3-3σ is YES-associated protein 1 (YAP1), which is part of the Hippo signaling pathway^[31]^. Phosphorylation of Ser127 induces binding to 14-3-3σ and inhibits YAP1 by preventing its nuclear localization^[32]^. We used the phosphorylated YAP1 peptide (residues 124-133) as a basis for the design of electrophile-containing peptides targeting Cys38 in 14-3-3σ. The top hit peptides irreversibly labeled 14-3-3σ within minutes. X-ray crystallography validated the predicted binding mode proposed by CovPepDock. Finally, the peptides label 14-3-3σ at nanomolar concentrations with exceptional selectivity in cell lysates.

## Results

### Method Overview

We implemented our covalent peptide docking protocol within the Rosetta modeling framework^[33,34]^ based on FlexPepDock^[23]^, to allow modeling of electrophilic peptides that covalently bind a receptor cysteine. Various electrophilic residues, modeled in their adduct form, were introduced to Rosetta, as well as a suitable reacted form of cysteine in the receptor protein which can covalently bind these electrophilic residues. The electrophilic groups may include non-natural or modified amino acids, capping groups and peptoids ^[35,36]^. A virtual atom is added to each residue, to represent the optimal placement of the atom at the other end of the covalent bond (Fig. 1A). During peptide docking, FlexPepDock iterates through several cycles of rigid-body optimization and flexible backbone sampling of the peptide^[23]^. During the full-atom refinement steps, we apply distance constraints between each of the covalent bond actual atoms and its virtual placeholder in the partnering residue. These constraints penalize the score of structures that diverge from the ideal bond length and angles. We parameterized 37 electrophilic side-chains, that can now be included in peptide docking simulations (Fig. 1B; Supplementary Fig. 1) Note, that due to the modular nature of Rosetta, these can now also be incorporated in any other Rosetta protocol such as protein-protein docking, structure prediction and design, and cyclic peptide design^[37]^.

**Figure 1.**
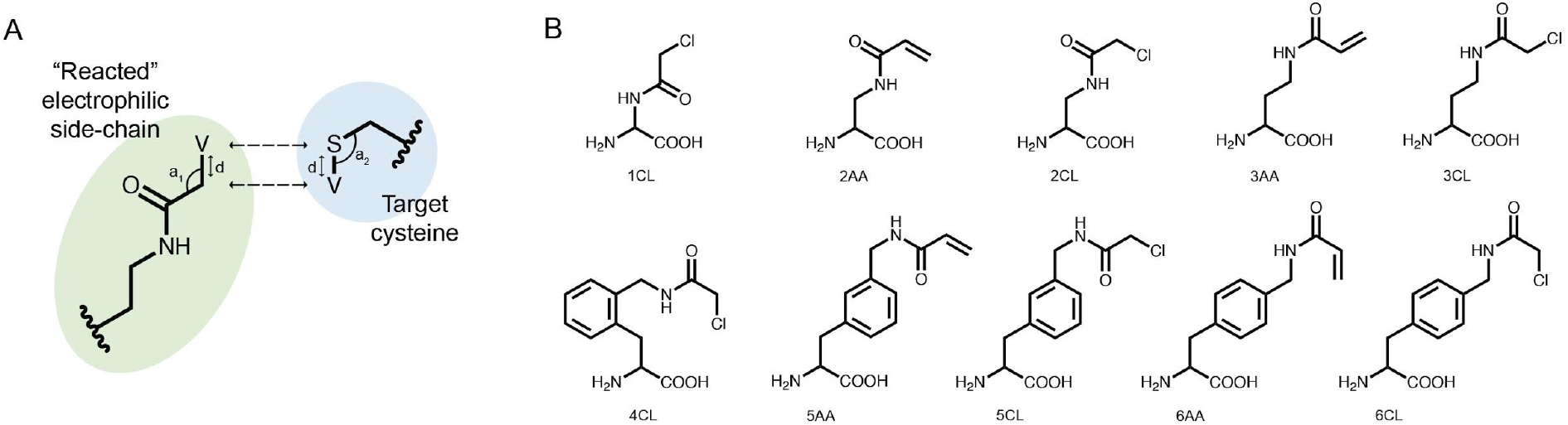
Implementing covalent binding in Rosetta. **A**. Virtual atoms (V) are added to both the electrophilic residue (green) and the cysteine on the target protein (blue). These define the optimal covalent bond length and angles (d, a_1_, a_2_), which are enforced through the addition of distance constraints between the actual atoms and the virtual placeholders (dashed arrows). **B**. Chemical structures of 10 out of 37 electrophilic side-chains parametrized for use in covalent docking. Note that these can be modeled in Rosetta as either L-or D-amino acids (See all of the implemented side-chains in Supplementary Fig. 1).

### CovPepDock accurately recapitulates known covalent peptide complexes

To assess the ability of our protocol to recapitulate known structures, we tested it against two benchmarks: a disulfides dataset (Supplementary dataset 1), composed of disulfide-mediated peptide-protein complexes; and an electrophiles dataset (Supplementary dataset 2), composed of peptide-protein complexes in which an electrophilic moiety on the peptide, such as a C-terminal aldehyde or ketone, N-terminal acrylamide and chloroacetamide caps, or a non-natural amino acid mediates a covalent bond to a receptor cysteine.

Following our previous criteria for the assessment of peptide docking^[23]^, we define two levels of success: a model is defined as near-native if the predicted peptide conformation is within 2Å interface backbone RMSD from the native conformation (interface residues are defined as any peptide residue whose Cβ atom is within 8Å of a Cβ atom of the binding partner, RMSD is calculated over all backbone atoms). Sub-angstrom models show interface backbone RMSD <1Å. The disulfides dataset comprises 115 structures, consisting of 32 different receptors in 49 unique complexes (Supplementary dataset 1). For 16 of the receptors in this dataset (24 complexes), a free, unbound structure was also available. We superimposed these structures onto their bound counterparts to create an unbound subset with the peptide positioned at the binding site. Disulfide-bonded cysteines have previously been implemented in Rosetta^[38]^. However, we created a new disulfide-bonded variant of cysteine to avoid any disulfide-specific behaviours or score terms encoded in Rosetta, that may affect our results. We docked each peptide starting from both its native conformation and an extended backbone conformation (where all the peptide φ/ψ dihedrals are set to +135°/ 135°, respectively). The latter simulates a more challenging scenario, where only the peptide sequence and approximate binding site is known.

We performed 100 docking simulations for each case and an additional 100 simulations with low-resolution pre-optimization for each extended peptide^[23]^. Our protocol sampled near-native models in 100% of the bound, non-extended cases, and ranked them among the top five interface-scoring models in 89% of the cases, and as the top scoring model in 78% (Fig. 2A). In the more difficult tasks of either unbound or extended docking, our protocol sampled near-native models in 87% and 43% of the cases, and ranked them in the top five scoring models in 83% and 20%, respectively. Under the more stringent criterion, CovPepDock ranked 64% and 37% sub-angstrom models of the bound and unbound sets respectively, in the top five scoring models (Fig. 2B; 45% and 25% as the top scoring model).

**Figure 2.**
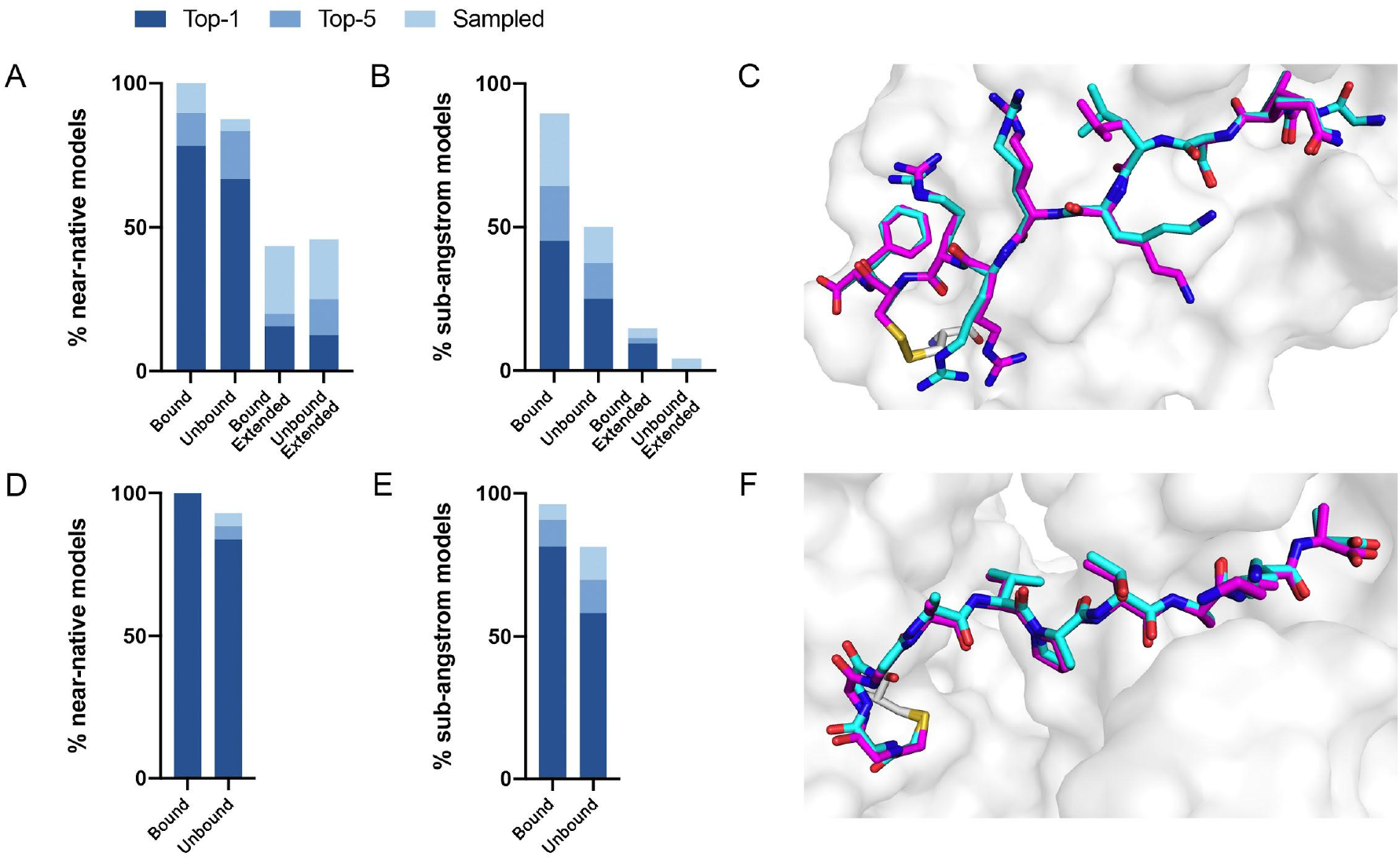
CovPepDock is successful in retrospective benchmarks. **A**. Percent of starting structures in the disulfides dataset, for which a near-native model (interface backbone RMSD < 2Å) was sampled (light blue), ranked among the top five scoring models (blue) or ranked as the top scoring model (dark blue). Similarly, **B** shows the percent of sub-angstrom models (interface backbone RMSD < 1Å) in the disulfides dataset. **C**. Native conformation (cyan) and top-scoring model (magenta) of PDB ID 5FGY from the disulfides dataset. **D**. and **E**. show the percent of near-native models and sub-angstrom models in the electrophiles dataset, accordingly, and **F**. shows the docking prediction for PDB ID 4WVI from the electrophiles dataset.

Similar docking studies were performed for the electrophiles dataset with 54 structures, consisting of 32 different receptors and 16 different electrophilic residues in 52 unique complexes (Supplementary dataset 2). An unbound structure was available for 24 receptors (43 complexes). Peptides in this dataset were modeled starting from their non-extended conformation only. Near-native models were sampled and ranked as the top interface-scoring model in 100% of the bound cases. In the unbound subset, near-native models were sampled and ranked among the top five interface-scoring models in 88% of the cases (Fig. 2D). Sub-angstrom models were ranked in the top five scoring models in 90% and 69% of the bound and unbound cases, respectively (Fig. 2E). The superior performance against the electrophile set might be explained by the different distribution of interface size between the datasets, where complexes in the electrophile set tend to have larger surface area buried at the interface (Supplementary Fig. 2).

### Design of Covalent Peptide Binders from Non-Covalent Templates

While there is much to be improved in the docking performance when starting from an extended peptide structure, in many practical cases, an initial peptide candidate is actually available. This can be either a known peptide binder or a protein fragment extracted from an interface with a binding partner. For instance, for protein-protein interactions, we can utilize continuous segments derived from the target interface to inhibit the parent interaction^[39–41]^. When an interfacial cysteine is available, modified, electrophilic versions of these peptides may result in superior inhibitors to the “wild-type” peptide/protein partner.

We developed a design protocol to identify the best peptide position for the incorporation of an electrophilic amino-acid and to select the optimal electrophilic side-chain. Starting from a peptide-protein complex template structure, we identify the peptide positions that are in close proximity to the target cysteine (Cα-Sγ distance < 10Å) and mutate each of these positions to various electrophilic amino acids. We then apply our covalent docking protocol to each of these putative complexes (Fig. 3). We introduced to Rosetta a set of 22 non-canonical electrophilic amino acids to be used within this protocol (Supplementary Fig. 1A; The rest of the 37 electrophiles were implemented to model complexes from the electrophilic peptides benchmark; see Supplementary Fig. 1B and Supplementary Dataset 2). We focused on acrylamide and chloroacetamide “warheads”, since these were shown to be sufficiently mild to ensure that the binding is driven mostly by recognition, rather than reactivity, thus reducing the number of off-targets in cellular applications^[42,43]^. All of the amino acid precursors in this set are commercially available as fmoc-derivatives, to encourage a broad use of our protocol in the future.

**Figure 3.**
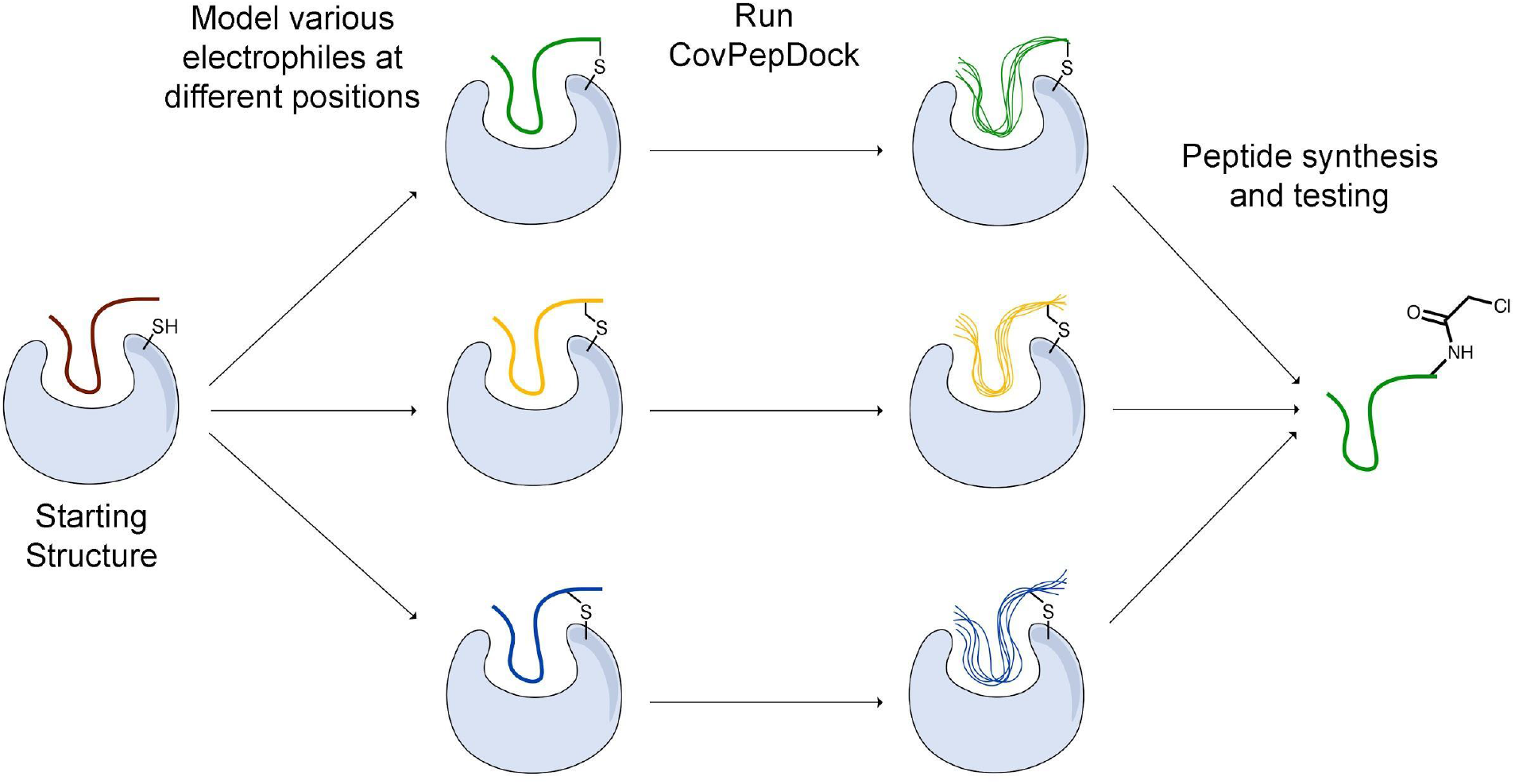
Outline of our covalent peptide design protocol. Starting from an initial peptide candidate that is at the interface with the target cysteine, we install various electrophiles in different peptide positions. We then use our docking protocol to model and evaluate their binding modes, and to select promising candidates for further testing and synthesis.

We applied this protocol to 49 of the disulfide-mediated complexes. For 23 out of the 32 protein targets, at least one electrophilic modification resulted in a top interface-scoring model with interface backbone RMSD < 1Å and constraint score < 2 (constraint score is a measure of the deviation of the covalent bond atoms from their virtual placeholder; the closer the score is to zero, the better the covalent bond fits the ideal geometry). These suggest that installing the electrophile does not interfere with the reversible binding, and is likely to result in a selective and potent irreversible peptide binder.

To evaluate the scope of targets to which this protocol can be applied, we searched the Protein Data Bank for cysteines that are at the interface with another protein chain. We used the Rosetta PeptiDerive application^[39]^ on each of these pairs, to identify short protein segments (3-15 amino acids) that have a significant contribution to the binding energy between the chains (interface score < 0), and that are in close proximity to the target cysteine (Cα-Sγ distance < 10Å). Our search yielded 7,154 protein pairs whose interfaces include such a segment, which can be used as an initial template for our protocol.

To make this protocol widely available we have implemented a free online version of the CovPepDock design protocol in ROSIE (Rosetta Online Server that Includes Everyone; https://rosie.graylab.jhu.edu/cov_pep_dock).

### CovPepDock identifies potent and selective 14-3-3σ covalent peptides

A potential target that was identified by our PDB-wide search is a cysteine residue at position 38 of the protein 14-3-3σ. We used CovPepDock prospectively to design covalent peptides to bind 14-3-3σ based on the non-covalent complex with YAP1 phosphopeptide (PDB ID: 3MHR)^[44]^. Amino-acids 131-133 of the YAP1 peptide were identified as potential sites for the installation of an electrophile (Supplementary Fig. 3), and each of these positions was mutated to the L-isomer of our 22 newly implemented electrophilic amino acids, as well as 8 D-isomers to which we had access, resulting in a total of 90 different designs (Supplementary dataset 3).

We synthesized seven peptides that received particularly high scores and low RMSD (peptides **1-7**), as well as three chloroacetamide-based peptides with bad scores and RMSD (Peptides **8-10**, Fig. 4A and Supplementary Fig. 4). To evaluate irreversible binding, we incubated the peptides with recombinant full-length 14-3-3σ and used intact protein liquid chromatography/mass spectrometry (LC/MS) to quantify the extent and rate of labeling of the protein. All top scoring peptides labeled the protein efficiently even with only a 2.5-fold excess (5 µM peptide/ 2 µM protein). As expected, acrylamide-based peptides were less reactive than chloroacetamides, reaching complete labeling of 14-3-3σ within several hours (Fig. 4B, Supplementary Fig. 5A) while some chloroacetamides labeled the protein fully within minutes (Figure 4C, Supplementary Fig. 5B). On average high-scoring peptides labeled 14-3-3σ more efficiently than low-scoring peptides, with one of the low scoring peptides showing no labeling even after extended incubation times. We also performed a dose response experiment conducted at constant incubation time of 5.5 hours (Supplementary Fig. 5C). This experiment implied that once a molar excess of peptide is reached, the labeling rate is independent of peptide concentration, suggesting that the non-covalent binding step is rapid and with high affinity and that the formation of the covalent bond within the noncovalent complex is the rate limiting step. To confirm this, we prepared a fluorescently labeled non-covalent analog of peptide **5** with a non-nucleophilic acetate group on position 133, and used fluorescence polarization to measure its binding affinity to 14-3-3σ (Supplementary Fig. 5D). We measured a dissociation constant (K_D_) of 106 ± 10 nM, indicating that in the conditions used in the covalent binding tests, the protein is fully saturated with bound peptide prior to covalent bond formation. To verify that the variation in labeling rates is not the result of difference in the intrinsic reactivity of the electrophiles, we performed a thiol reactivity assay^[45]^ using reduced DTNB (Ellman’s reagent) as the nucleophile (Supplementary Fig. 6). All chloroacetamide-containing peptides displayed similar reaction rates, while acrylamide-containing peptides showed no reaction with DTNB. These results indicate that the observed labeling of 14-3-3σ is indeed due to recognition and favored geometry of the electrophile within the bound complex.

**Figure 4.**
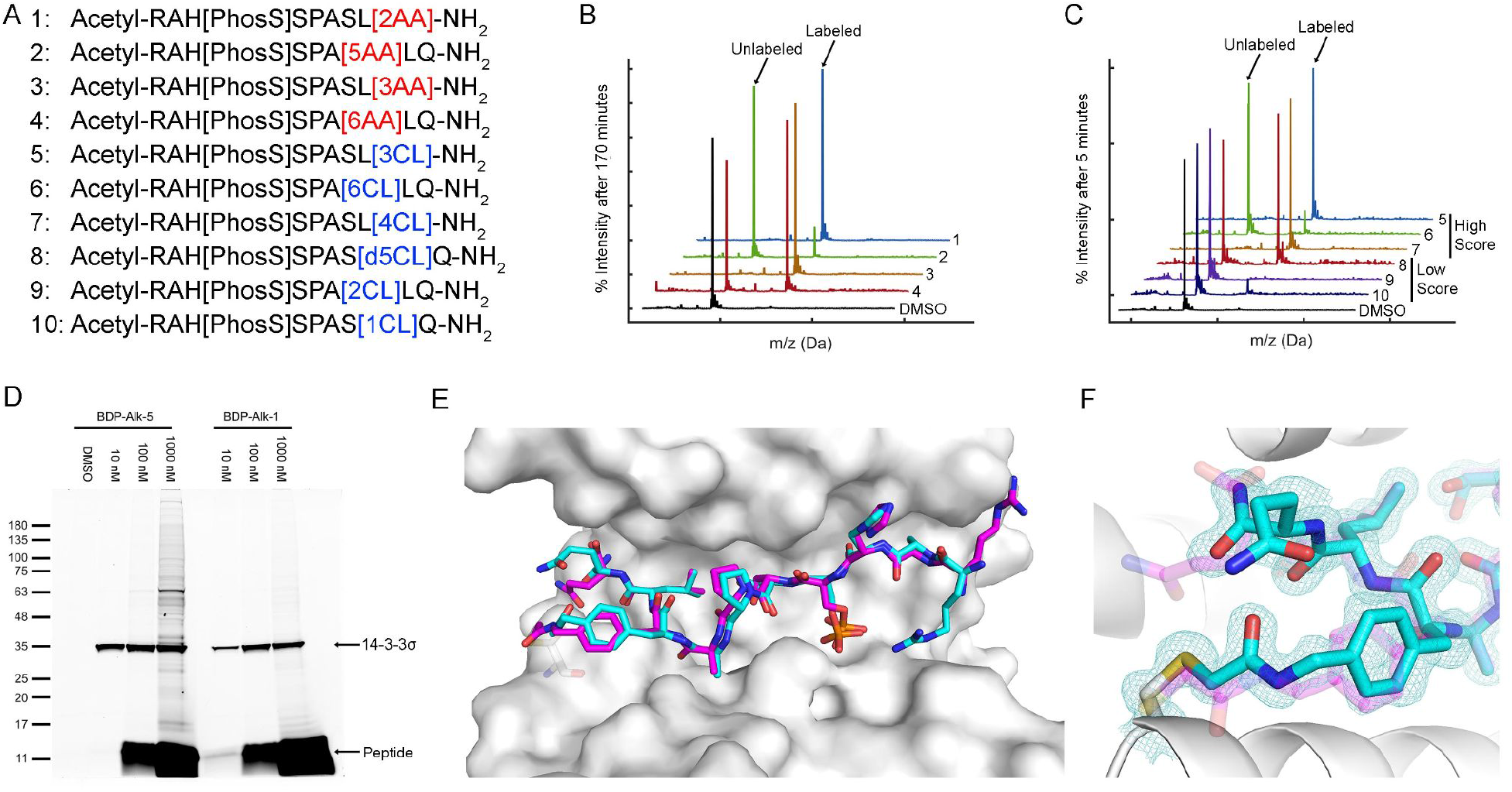
CovPepDock identifies potent and selective irreversible 14-3-3σ binders. **A**. List of candidate binding peptides for 14-3-3σ. AA indicates acrylamide, CL indicates chloroacetamides (see amino-acid structures in Fig.1B). **B**. Intact LCMS monitoring of the reaction of 14-3-3σ with the acrylamide peptides 1-4, after 170 minutes of incubation at 5 µM peptide/ 2 µM protein. **C** Intact LCMS monitoring of the reaction of 14-3-3σ with the chloroacetamide peptides 5-10, after 5 minutes of incubation at 5 µM peptide/ 2 µM protein. **D**. Reaction of fluorescently labeled electrophilic derivatives of peptides **1** and **5** with 14-3-3σ in A431 cell lysates, separated on a 4-20% Bis Tris SDS gel. Fluorescence was measured using a 532 nm excitation laser and a 550 nm longpass filter. **E**. Structural overlay of the covalent docking prediction of peptide 6 (magenta sticks) and the co-crystal structure of peptide **6** (cyan sticks) when bound to 14-3-3σ (white surface). **F**. Close-up view of the electrophilic residue in peptide **6**, including its electron density. 2F_o_-F_c_ electron density maps are contoured at 1σ (PDB: 7O07).

To test the ability of the electrophilic peptides to bind 14-3-3σ in the context of the cellular proteome, we synthesized fluorescently labeled derivatives of the most potent acrylamide peptide **1** and most potent chloroacetamide peptide **5**, and incubated them with A431 cell lysates. The peptides displayed potent and highly selective labeling of 14-3-3σ in the cell lysates at concentrations as low as 10 nM (Fig. 4D). For peptide **5**, the binding was so efficient that at 10 nM peptide, the added peptide was entirely consumed by 14-3-3σ in the lysate without any free peptide remaining and without significant off-target binding. At higher peptide concentrations off-target binding was observed, particularly for the more intrinsically reactive chloroacetamide **5**. To confirm that the observed target was indeed 14-3-3σ, we also performed western blot following incubation with the lysate with the fluorescent analog of **5** (supplementary Fig. 7). We detected 14-3-3σ using an antibody raised against residues 25-38, an epitope that contains the target cysteine (cys38) and is not shared with other 14-3-3 members. Incubation with the peptide resulted in replacement of the 14-3-3σ band with a slightly higher molecular weight band with considerably lower intensity, indicating impaired recognition of the epitope by the antibody due to peptide binding. The new band was confirmed to contain the peptide via fluorescence. These results confirm that the peptide reacted with 14-3-3σ potently and selectively in cell lysates.

To validate the predicted binding mode, we crystallized the electrophilic peptides with recombinant 14-3-3σ. Most peptides gave either no crystals or insufficient density to determine the conformation of the electrophilic residues. However, peptide **6** provided a high resolution dataset with unambiguous density for the electrophilic residue and the covalent bond, confirming the covalent binding to Cys38 in 14-3-3σ (figure 4E,F). The predicted structure showed the closest agreement to the obtained structure along the inner backbone of the peptide (interface backbone RMSD = 0.9Å), with some differences in the edges, such as an opposite orientation of the N-terminal arginine and a flip in the direction of the amide bond nearest to the electrophile. This close agreement is in line with our retrospective benchmarks and highlights the accuracy of CovPepDock.

## Discussion

Covalent peptides offer new opportunities to bind very challenging targets that are currently considered undruggable. In this study, we developed a pipeline that extends Rosetta FlexPepDock to enable modeling of electrophilic peptides that form a covalent bond to a receptor cysteine. Benchmarking this method against available structures of peptide-protein complexes showed high accuracy. On the basis of this method, we developed a design protocol that creates electrophilic peptides based on a template peptide-protein complex structure. This protocol can be useful for discovering novel covalent peptide binders for inhibiting protein-protein or peptide-protein interactions, as well as for enhancing the potency and selectivity of previously identified peptide binders. The protocol is freely available as part of the ROSIE web-server (https://rosie.graylab.jhu.edu/cov_pep_dock).

An important feature of our protocol is that it can easily incorporate a wide variety of structural modifications included in the Rosetta database, thus allowing modeling and design of peptides that far surpass the 20 natural L-amino-acid space. Examples include non-natural, N-methylated or D-amino acids, terminal caps and peptoids. Such modifications have the potential to improve the pharmacokinetic properties of the peptide, such as bioavailability and resistance to proteolytic degradation, which is one of the major challenges in designing peptide-based drugs. Moreover, the steps used in our protocol to achieve peptide-protein cross-chain covalent binding, can be adapted to create intra-chain covalent linkages, such as peptide cyclizations, helix stapling and hydrogen bond surrogates (HBS), which have been shown to enhance membrane permeability and proteolytic stability^[46–50]^. These modifications also stabilize the desired conformation of the peptide, thus reducing the conformational entropy and increasing the binding affinity. In addition we are currently working towards incorporation of these new electrophiles into peptoids^[35,36]^.

As an extension of FlexPepDock, our protocol inherits a few limitations. FlexPepDock incorporates full flexibility for the receptor and peptide side-chains, as well as for the peptide backbone, but lacks any flexibility for the receptor backbone. Manual inspection of complexes for which a near-native model was sampled in the bound docking, but not in the unbound docking, showed that in all six cases, a slight movement of the receptor backbone is crucial for binding. For example, the backbone of the unbound receptor in PDB: 6B9Z clashes with the peptide in the bound PDB: 6B9Y. In another example, the receptor cysteine in the unbound PDB: 5LAR is shifted 11.3Å away from the peptide in the bound PDB: 3P4K, and therefore cannot form the disulfide bond. Additionally, the decrease in success rate in the extended docking mode, implies that a more extensive sampling method, such as the fragment-based FlexPepDock *ab initio* protocol^[51]^, might be necessary to improve the performance of our protocol.

Nevertheless, we were able to successfully apply CovPepDock to design covalent peptide binders for 14-3-3σ. To our knowledge these are the first covalent peptide binders for this target. All predicted binders were able to covalently label 14-3-3σ, with two chloroacetamides (**5** and **7**) and two acrylamides (**1** and **3**) particularly rapidly. Covalent binding is a two step process, in the first a reversible encounter complex is formed, followed by irreversible binding. Unless the new electrophilic residue introduces a major clash, we assume the rate of the first step is similar for all peptides, and have measured the K_D_ for a non-covalent analog to be around 100 nM (Supplementary Fig. 5D). Two factors can accelerate the rate of the second step. First is intrinsic thiol reactivity, which is higher for chloroacetamides (Supplementary Fig. 6) and explains their faster binding. Second is exact orientation of the electrophile with respect to the nucleophile. This likely explains the differences between the labeling rates. As a stringent test we chose three high-reactivity chloroacetamide peptides that did not rank well by docking. One of them was not able to label at all, and the other two labeled slower than the high-scoring predictions.

The utility of these potent 14-3-3σ covalent binders was demonstrated by their ability to very selectively label the protein in a cellular lysate (Fig. 4D), a task that might prove difficult even for antibodies, due to the high conservation of 14-3-3 proteins. The 14-3-3σ isoform plays a unique role amongst 14-3-3 proteins and this reagent might enable to shed more light on its biology. Future optimization of the sequence with cell penetrating tags may enable selective and potent competitive inhibition of 14-3-3σ based on these binders.

We have identified thousands of possible proteins that may be amenable to covalent binding by this approach. We have made the design protocol available through a free web-server and based it on commercially available amino-acids. We hope this would facilitate broad usage to generate a wide range of covalent peptides for chemical biology and potentially drug discovery applications.

## Methods

### Benchmarks

To construct the disulfides dataset, we searched the PDB using the following attributes: (1) the experimental method is X-ray diffraction, (2) the structure contains at least two protein entities, (3) it contains a chain with sequence length between 3 and 15, and (4) it contains at least one disulfide bond. We then filtered the results for structures that contain a disulfide bond between a cysteine residue on a peptide chain (3-15 amino acids long) and a cysteine residue on a non-peptide chain.

Similarly, to construct the electrophiles dataset, we searched the PDB using attributes (1)-(3) and (5) the structure contains a covalent link from a cysteine SG atom. We filtered the results for structures that contain a link between a non-cysteine residue on a peptide chain and a cysteine residue on a non-peptide chain, and that do not contain other non-canonical peptide residues other than the electrophilic linkage.

The unbound structures were collected using the PDB sequence similarity search, with identity cutoff of 95%.

The receptors in both datasets were clustered using the PDB sequence clusters with 70% sequence identity. The peptides were clustered manually by 100% sequence and length identity.

### Parametrizing New Residues

The non-canonical amino acids used in this study were implemented in Rosetta using the protocol described in Renfrew et al.^[35]^. We first use the GaussView^[52]^ interface to draw the residue in its reacted form, with an acetylated N-terminus, a methylamidated C-terminus, and either a methylthiolated side-chain in case of an electrophilic residue, or an ethylated thiol in case of the electrophile-binding variant of cysteine. We then optimize the structure using the Gaussian software^[53]^ with the following options: HF/6-31G(d) scf=tight test. We convert the optimized structure to a mol file using OpenBabel toolbox^[54]^ (http://openbabel.org), and then to a Rosetta residue ‘params file’ using the molfile_to_params_polymer.py script provided in Rosetta. Rotamer libraries are generated using the Rosetta MakeRotLib application.

To allow a residue to covalently bind another residue, we use similar steps to these described in Drew et al.^[36]^ for oligooxopiperazines. We add a CONNECT record to the residue params file, specifying which atom participates in the inter-residue covalent bond. We also add a virtual atom, and define its internal coordinates according to the optimal position of the atom at the other end of the covalent bond, as predicted by the Gaussian optimization. This virtual atom is used during docking to favor the correct covalent bond geometry.

Chiral amino acids were implemented as L-isomers. D-isomers were modeled using the D_AA.txt patch file provided in the Rosetta database.

### Covalent Docking

Our protocol follows the FlexPepDock workflow described in Raveh et al.^[23]^. We start by prepacking the input structure, and then use the prepacked structure as a starting point for the refinement step, in which we generate 100 models of the complex. Command-line templates are provided below. For docking of extended peptides, we perform an additional 100 refinement runs with a low-resolution pre-optimization step, using the options shown in parenthesis.

To ensure proper sampling and scoring of the covalent bond during the refinement step, we apply AtomPair constraints between each of the covalent bond atoms and its virtual placeholder in the partnering residue. We use the HARMONIC score function, centered at 0 and with a standard deviation of 0.3. During the low-resolution pre-optimization step, we impose an AtomPair constraint between the Cα atoms of the two bonded residues, using the FLAT_HARMONIC score function, centered at 0, with a tolerance equals to the sum of the length of the bonds connecting these atoms, and with a standard deviation of 0.1.

Prepacking command: ROSETTA_BIN/FlexPepDocking.linuxgccrelease -s START.pdb -native NATIVE.pdb -flexpep_prepack -ex1 -ex2aro -extra_res_fa NCAA.params -receptor_chain REC_CHAINS -flexPepDocking:peptide_chain PEP_CHAIN (-extend_peptide) Refinement command: ROSETTA_BIN/FlexPepDocking.linuxgccrelease -s PREPACKED.pdb -native NATIVE.pdb -pep_refine -ex1 -ex2 -use_input_sc -extra_res_fa NCAA.params -cst_fa_file FA_CST -cst_fa_weight 1 -receptor_chain REC_CHAINS -flexPepDocking:peptide_chain PEP_CHAIN (-extra_res_cen NCAA.cen.params -constraints:cst_file CEN_CST -cst_weight 1 -lowres_preoptimize)

### PDB-wide search for potential targets

We searched the PDB for X-ray crystal structures that contain at least two protein entities. We then searched each of these structures for cysteine residues that are at the interface with another protein chain (Cα-Sγ distance < 10Å). For each of these chain pairs, we used Rosetta PeptiDerive^[39]^ to extract protein fragments of 3-15 amino acids from the interface, with the cysteine-containing chain acting as the receptor and the other chain as the partner protein. Finally, we filtered the results for derived peptides with interface score < 0, and that have at least one Cα atom within 10Å from the Sγ atom of a receptor cysteine.

We clustered the results using the PDB sequence clusters, with 90% sequence identity for both the receptor chain and the partner chain (we ignored cases in which one of the chains did not belong to any cluster). This resulted in 7,154 protein pairs whose interface contain a potential target cysteine and an initial peptide candidate.

### 14-3-3σ peptide design

We used PDB: 3MHR as a template structure in our design protocol. We used Rosetta fixed backbone design application (fixbb) to mutate positions 131-133 of the peptide to the L-isomer of amino acids 1-11AA and 1-11CL, and to the D-isomer of amino acids 4-6AA, 4-6CL, 9AA and 9CL. We then applied CovPepDock to generate 100 models of each of these mutated peptides, using the prepacking and refinement command-lines shown in the “Covalent Docking” method section.

We manually inspected the 900 top-interface-scoring models, focusing on near-native models with constraint score < 2. We selected 7 high-ranking peptides that vary in their mutated position, geometry and reactivity. To select the 3 low-ranking peptides, we inspected the 13 designs for which all the top 10 interface-scoring models had interface backbone RMSD > 2Å.

### Peptide Synthesis

Reagents for peptide synthesis were purchased from Chem-Impex. Peptides were synthesized on Rink Amide resin using standard fmoc chemistry on a 0.025 mmol scale. The resin was swelled for 30 minutes in dichloromethane (DCM), then washed with dimethylformamide (DMF). Fmoc deprotections were carried out using 20% piperidine in DMF (3×3 minutes), and couplings were performed as follows: 4 equivalents of amino acid were mixed with 4 equivalents of HATU and 8 equivalents of DIPEA in DMF and added to the resin with mixing for 45 minutes. For phosphoserine, propargylglycine and amino acids used for introducing the electrophile, 2 equivalents were used and reaction times were extended to 2 hours. After the last fmoc deprotection, the peptides were acetylated at the N terminus using acetic anhydride (10 equivalents) and DIPEA (20 equivalents) in DMF for 30 minutes. Finally the resins were washed with DCM, dried in a dessicator, and cleaved using 85% TFA/5% thioanisole/5% ethanedithiol/2.5% TIPS/2.5% water for 2 hours with tumbling. The cleaved peptides were precipitated in cold diethyl ether, washed once with ether, dried, dissolved in 50% acetonitrile and lyophilized.

The electrophile was introduced to the peptides via various amino acids containing boc-protected amines. Since the peptides are N-terminally acetylated and contain no lysines, the amine was expected to be acylated readily and selectively with acrylic acid, chloroacetic acid or acetic acid. To acylate the peptides, the crude peptide was dissolved in DMF to an estimated concentration of 0.1 M. The acid to be coupled was dissolved in DMF at 0.2 M, and an activation mixture of 0.11 M EDC/0.11M HOBT/0.5M DIPEA in DMF was prepared as well. 75 µl of activation mixture was mixed with 75 µl of acid solution, generating the activated acid with no excess of EDC. After 5 minutes this was added to 75 µl of peptide solution, and mixed for 2 hours. The coupled peptide was diluted with 20% acetonitrile/water + 0.1% TFA and purified by HPLC. This yielded 0.5-6 mg of pure peptide for the acylated peptides, with chloroacetamides and acetates giving higher yields than acrylamides. Following HPLC the purified fractions were lyophilized from 30% acetic acid to remove the TFA, dissolved in DMSO and stored at −80°C.

To prepare fluorescently labeled peptides, a residue of propargylglycine was coupled to the peptide at the N-terminus prior to N-terminal acetylation, cleavage and acylation with the electrophile. The pure peptide was then labeled as follows using copper-catalyzed azide-alkyne cycloaddition (CuAAC): 4 µl of 20 mM alkyne-peptide in DMSO were mixed with 10 µl of 5 mM BDP-TMR-azide (Lumiprobe). To the mixture was added 50 µl water, 0.8 µl of 0.1 M CuSO4 : Tris(3-hydroxypropyltriazolylmethyl)amine, 30 µl tert-butanol, and 2 µl of 80 mM freshly dissolved sodium ascorbate. The reaction proceeded in the dark at room temperature for 30 minutes, and another 2 µl of 80 mM sodium ascorbate was added. After 30 minutes the labeled peptide was purified by HPLC.

All peptides were characterized by LC-MS (Supplementary dataset 4).

### Protein expression and purification used for MS and FP experiments

#### 14-3-3s full length (1-248)

*<u>SYYHHHHHHDYDIPTTENLYFQGAMGS</u>MERASLIQKAKLAEQAERYEDMAAFMKGAVEKGEE LSCEERNLLSVAYKNVVGGQRAAWRVLSSIEQKSNEEGSEEKGPEVREYREKVETELQGVCDTVLGLLDSHLIKEAGDAESRVFYLKMKGDYYRYLAEVATGDDKKRIIDSARSAYQEAMDISKKEMPPTNPIRLGLALNFSVFHYEIANSPEEAISLAKTTFDEAMADLHTLSEDSYKDSTLIMQLLRDNLTLWTADNAGEEGGEAPQEPQS*

#### 14-3-3s DC (1-231)

*<u>SYYHHHHHHDYDIPTTENLYFQ|GAMGS</u>MERASLIQKAKLAEQAERYEDMAAFMKGAVEKGEE LSCEERNLLSVAYKNVVGGQRAAWRVLSSIEQKSNEEGSEEKGPEVREYREKVETELQGVCDTVLGLLDSHLIKEAGDAESRVFYLKMKGDYYRYLAEVATGDDKKRIIDSARSAYQEAMDISKKEMPPTNPIRLGLALNFSVFHYEIANSPEEAISLAKTTFDEAMADLHTLSEDSYKDSTLIMQLLRDNL TLWT*

*C-terminal truncation is made to improve crystallization of 14-3-3*.

*<u>.....</u> = purification tag*

| *= TEV cleavage site*

A pPROEX HTb expression vector encoding the human 14-3-3σ with an N-terminal his6-tag was transformed by heat shock into NiCo21 (DE3) competent cells. Single colonies were cultured in 50 mL LB medium (10 mg/mL ampicillin). After overnight incubation at 37 °C, cultures were transferred to 2 L TB media (10 mg/mL ampicillin, 1 mM MgCl2) and incubated at 37 °C until an OD600 nm of 0.8-1.2 was reached. Protein expression was then induced with 0.4 mM isopropyl-β-d-thiogalactoside (IPTG), and cultures were incubated overnight at 18°C. Cells were harvested by centrifugation (8600 rpm, 20 minutes, 4 °C) and resuspended in lysis buffer (50 mM Hepes, pH 8.0, 300 mM NaCl, 12.5 mM imidazole, 5 mM MgCl2, 2 mM βME) containing cOmplete™ EDTA-free Protease Inhibitor Cocktail tablets (1 tablet/ 100 ml lysate) and benzonase (1 μl/ 100 ml). After lysis using a C3 Emulsiflex-C3 homogenizer (Avestin), the cell lysate was cleared by centrifugation (20000 rpm, 30 minutes, 4 °C) and purified using Ni2+-affinity chromatography (Ni-NTA superflow cartridges, Qiagen). Typically two 5 mL columns (flow 5 mL/min) were used for a 2 L culture in which the lysate was loaded on the column washed with 10 CV wash buffer (50 mM Hepes, pH 8.0, 300 mM NaCl, 25 mM imidazole, 2 mM βME) and eluted with several fractions (2-4 CV) of elution buffer (50 mM Hepes, pH 8.0, 300 mM NaCl, 250 mM imidazole, 2 mM bME). Fractions containing the 14-3-3σ protein were combined and dialyzed into 25 mM HEPES pH 8.0, 100 mM NaCl, 10 mM MgCl2, 500 μM TCEP. Finally, the protein was concentrated to ∼60 mg/mL, analyzed for purity by SDS-PAGE and Q-Tof LC/MS and aliquots flash-frozen for storage at −80°C.

### Protein expression and purification used for crystallography

A pPROEX HTb expression vector encoding the human 14-3-3σ truncated after T231 14-3-3σ Δc) and with an N-terminal his6-tag was transformed by heat shock into NiCo21 (DE3) competent cells. Single colonies were cultured in 50 mL LB medium (10 mg/mL ampicillin). After overnight incubation at 37 °C, cultures were transferred to 2 L TB media (10 mg/mL ampicillin, 1 mM MgCl2) and incubated at 37 °C until an OD600 nm of 0.8-1.2 was reached. Protein expression was then induced with 0.4 mM isopropyl-β-d-thiogalactoside (IPTG), and cultures were incubated overnight at 18°C. Cells were harvested by centrifugation (8600 rpm, 20 minutes, 4 °C) and resuspended in lysis buffer (50 mM Hepes, pH 8.0, 300 mM NaCl, 12.5 mM imidazole, 5 mM MgCl2, 2 mM βME) containing cOmplete™ EDTA-free Protease Inhibitor Cocktail tablets (1 tablet/ 100 ml lysate) and benzonase (1 μl/ 100 ml). After lysis using a C3 Emulsiflex-C3 homogenizer (Avestin), the cell lysate was cleared by centrifugation (20000 rpm, 30 minutes, 4 °C) and purified using Ni2+-affinity chromatography (Ni-NTA superflow cartridges, Qiagen). Typically two 5 mL columns were used for a 2 L culture in which the lysate was loaded on the column washed with 10 CV wash buffer (50 mM Hepes, pH 8.0, 300 mM NaCl, 25 mM imidazole, 2 mM βME) and eluted with several fractions (2-4 CV) of elution buffer (50 mM Hepes, pH 8.0, 300 mM NaCl, 250 mM imidazole, 2 mM βME). Fractions containing the 14-3-3σ protein were combined and dialyzed into 25 mM HEPES pH 8.0, 200 mM NaCl, 10 mM MgCl2, 2 mM BME. In addition, 1 mg TEV was added for each 100 mg purified protein to remove the purification tag. The cleaved sample was then again loaded on a 10 mL Ni-NTA column to separate the cleaved product from the expression tag and residual uncleaved protein. The flowthrough was loaded on a Superdex 75 pg 16/60 size exclusion column (GE Life Sciences) using 25 mM HEPES, 100 mM NaCl, 10 mM MgCl2, 500 μM TCEP (adjusted to pH=8.0) as running buffer. Fractions containing the 14-3-3σ protein were pooled and concentrated to ∼60 mg/mL, analyzed for purity by SDS-PAGE and Q-Tof LC/MS and aliquots flash-frozen for storage at −80 °C.

#### 14-3-3σ crystallography

Peptide and 14-3-3σ protein were dissolved in complexation buffer (20 mM HEPES pH 7.5, 100 mM NaCl, 10 mM MgCl2 and 20 μM TCEP) using a final 14-3-3σ concentration of 10 mg/mL and 12.5 mg/mL and a 1:1.2 and 1:2 molar stoichiometry of protein:peptide. These complexes were incubated overnight at room temperature. After the incubation, sitting-drop crystallization plates were set up in which each of the four complexation mixtures was combined with 24 crystallization buffers, optimized for 14-3-3σ crystallization (0.095 M HEPES (pH7.1, 7.3, 7.5, 7.7), 0.19 M CaCl2, 24, 25, 26, 27, 28 29 % (v/v) PEG 400 and 5% (v/v) glycerol). Herein a 1:1 mix (both 250 nL) of complexation mixture and crystallization buffer was made for crystal growth. Crystals grew within 10-14 days at 4 °C.

Suitable crystals were fished and flash-cooled in liquid nitrogen. X-ray diffraction (XRD) data were collected either at the European Synchotron Radiation Facility (ESRF) beamline ID30B, Grenoble, France, equipped with a PILATUS3 6M detector. Typical setting were 1440 image, 0.25°/image, 3% transmission and 0.1 s exposure time.

Data was processed using the CCP4i2 suite (version 7.1.10)^[55]^. DIALS^[56]^ was used to index and integrate the data after which scaling was done using AIMLESS^[57,58]^. The data was phased with MolRep (20057045), using protein data bank (PDB) entry 5N75 as a template. Ligand restraints for non-natural amino acids were generated with AceDRG^[59]^. Sequential model building (based on visual inspection Fo-Fc and 2Fo-Fc electron density map) and refinement were performed with COOT and REFMAC, respectively^[60–62]^. Finally, alternating cycles of model improvement and refinements were performed using coot and phenix.refine from the Phenix software suite (version 1.15.2-3472)^[63,64]^. Pymol (version 2.2.3) was used to make the figures and the structures were deposited in the protein data bank (PDB) with ID: 7O07. See table S2 for crystal statistics.

### LCMS instrumentation and runs

The LC/MS runs for 14-3-3σ were performed on a Waters ACQUITY UPLC class H instrument, in positive ion mode using electrospray ionization. UPLC separation used a C4-BEH column (300 Å, 1.7 μm, 21 mm × 100 mm). The column was held at 40 °C and the autosampler at 10 °C. Mobile phase A was 0.1% formic acid in water, and mobile phase B was 0.1% formic acid in acetonitrile. The run flow was 0.4 mL/min. The gradient used was 20% B for 2 min, increasing linearly to 80% B for 2.5 min, holding at 80% B for 0.5 min, changing to 20% B in 0.2 min, and holding at 20% for 0.8 min. The MS data were collected on a Waters SQD2 detector with an m/z range of 2–3071.98 at a range of 900–1900 m/z. The desolvation temperature was 500 °C with a flow rate of 800 L/h. The voltages used were 1.00 kV for the capillary and 24 V for the cone. Raw data were processed using openLYNX and deconvoluted using MaxEnt with a range of 20000:40000 Dat and a resolution of 1.5 Da/channel.

The LS/MS runs for peptides were performed using the same instrument with a C18-CSH column (300 Å, 1.7 μm, 21 mm × 100 mm) using a gradient starting from 1% B for 1 minutes, rising to 90% B in 4.5 minutes, holding at 90% B for 0.75 minutes, then decreasing to 1% B in 0.75 minutes and holding at 1% B for 1 minute. MS data were collected at a range of 80-2500 m/z, using identical conditions for ionization as with the protein.

### Intact LCMS monitoring of 14-3-3σ binding by the peptides

Recombinant 14-3-3σ was diluted to 2 µM in 25 mM HEPES pH 7.5, 100 mM NaCl, 10 mM MgCl2 and 20 µM TCEP, and the peptides were diluted to 100X the final concentration in DMSO. Labeling was initiated by diluting the peptide 100-fold into the protein solution and incubating the mixture at room temperature. The reaction was stopped at defined times by adding 6 µl of 2.4% formic acid to 24 µl of sample and immediately injecting 10 µl into for intact LCMS analysis.

### Binding of fluorescent electrophilic peptides to 14-3-3σ in A431 lysates

A-431 cells were grown at 37°C in a 5% CO2 humidified incubator and cultured in DMEM supplemented with 10% Fetal bovine serum,1% L-Glutamine, 1% Sodium Pyruvate and 1% pen-strep solution (all from Biological Industries).

A431 cell pellets were dispersed in lysis buffer (Tris 50 mM pH = 8, 150 mM NaCl, 1% IGEPAL + protease inhibitor cocktail P8340 from Sigma) and incubated on ice with occasional vortexing for 15 minutes. The cells were then centrifuged at 21000 g for 10 minutes at 4 °C, and the supernatant was collected. The total protein concentration was estimated using BCA protein assay (Pierce) and samples containing 40 µg protein in 20 µl buffer were incubated with fluorescent peptides for 90 minutes at room temperature in the dark. The sample were then denatured using NuPage LDS buffer + 20 mM DTT at 70 °C for 10 minutes, and loaded on 4-20% Bis-Tris gradient gels. After electrophoresis the gels were fixed (using two portions of 45% methanol/10% acetic acid/45% water for 10 minutes each) and then immersed in 100 mM Tris pH=8 in water and imaged using Typhoon using a 532 nm laser. ImageJ was used to process the images.

For western blot experiment, the gel was transferred to a nitrocellulose membrane using TransBlot Turbo (Bio Rad), and blocked with 5% BSA in TBST for 1 hour RT. The membrane was incubated with mouse anti 14-3-3σ antibody (Santa Cruz, sc-166473, diluted 1:200 in 5% BSA/TBST) overnight at 4 °C with gentle shaking. The membrane was then washed 3 times with TBST (5 minutes) and incubated with anti-mouse HRP antibody (CST 7076, diluted 1:2000 in 5% BSA/TBST) for 1 hour RT. The gel was washed again with TBST, and imaged using electrochemluminescence using GelDoc, followed by measurement of fluorescence using a 532 nm laser using Typhoon.

### Intrinsic reactivity measurement of electrophilic peptides

The reactivity of the electrophilic peptides was estimated using a DTNB assay^[45]^. Solutions of 20 mM peptide in DMSO were diluted to 400 µM in NaPi 25 mM pH = 7.4, 150 mM NaCl, and 25 µl of each peptide was placed in wells in a 384 well black, flat and transparent bottom plate. Then, 100 µM of DTNB was incubated for 10 minutes with 400 µM TCEP in the same buffer, at room temperature, to generate reduced DTNB. Reaction was initiated by adding 25 µl reduced DTNB solution to each peptide well, and monitoring the absorbance at 412 nm every 15 minutes at 37 °C with shaking, using a Tecan Spark plate reader.. As a blank, the absorbance of the same samples without added DTNB was measured and subtracted. Measurements were performed in triplicates.

### Fluorescence Polarization binding studies

Fluorescence polarization experiments were performed in 25 mM HEPES pH 7.5, 100 mM NaCl, 10 mM MgCl2 and 20 µM TCEP, with the addition of 0.05% IGEPAL to prevent nonspecific adsorption of the protein to the well surface. 5 nM fluorescent peptide was incubated with increasing concentrations of 14-3-3σ and the fluorescence polarization was measured at 37 °C using a Tecan Spark plate reader. Excitation was performed using a 550 ± 10 nm filter and emission was measured using a 620 ± 20 nm filter.

## Author Contributions

B.T., R.G and N.L. wrote the manuscript with input from all authors; B.T. designed and implemented the protocol, performed all computational work and peptide design; R.G. synthesized peptides, tested their activity; B.A.S expressed recombinant protein and performed X-ray crystallography; P.J.C aided in experimental design and peptide binding studies; C.O. oversaw and led all 14-3-3 related experiments; N.L. conceived and led the study.

## Data availability

CovPepDock protocol is freely available to use as part of the ROSIE web-server (https://rosie.graylab.jhu.edu/cov_pep_dock); Crystal structures described in this manuscript have been deposited to PDB (7O07); Additional experimental data is available in the extended data files.

## Acknowledgments

N.L. is the incumbent of the Alan and Laraine Fischer Career Development Chair. N.L. thanks the Israel Science Foundation for funding (grant no. 2462/19), The Israel Cancer Research Fund, and the Moross integrated cancer center. N.L. is also supported by the Helen and Martin Kimmel Center for Molecular Design, Joel and Mady Dukler Fund for Cancer Research, the Estate of Emile Mimran and Virgin JustGiving and the George Schwartzman Fund.

