## Supplementary Information for "Covalent Flexible Peptide Docking in Rosetta"

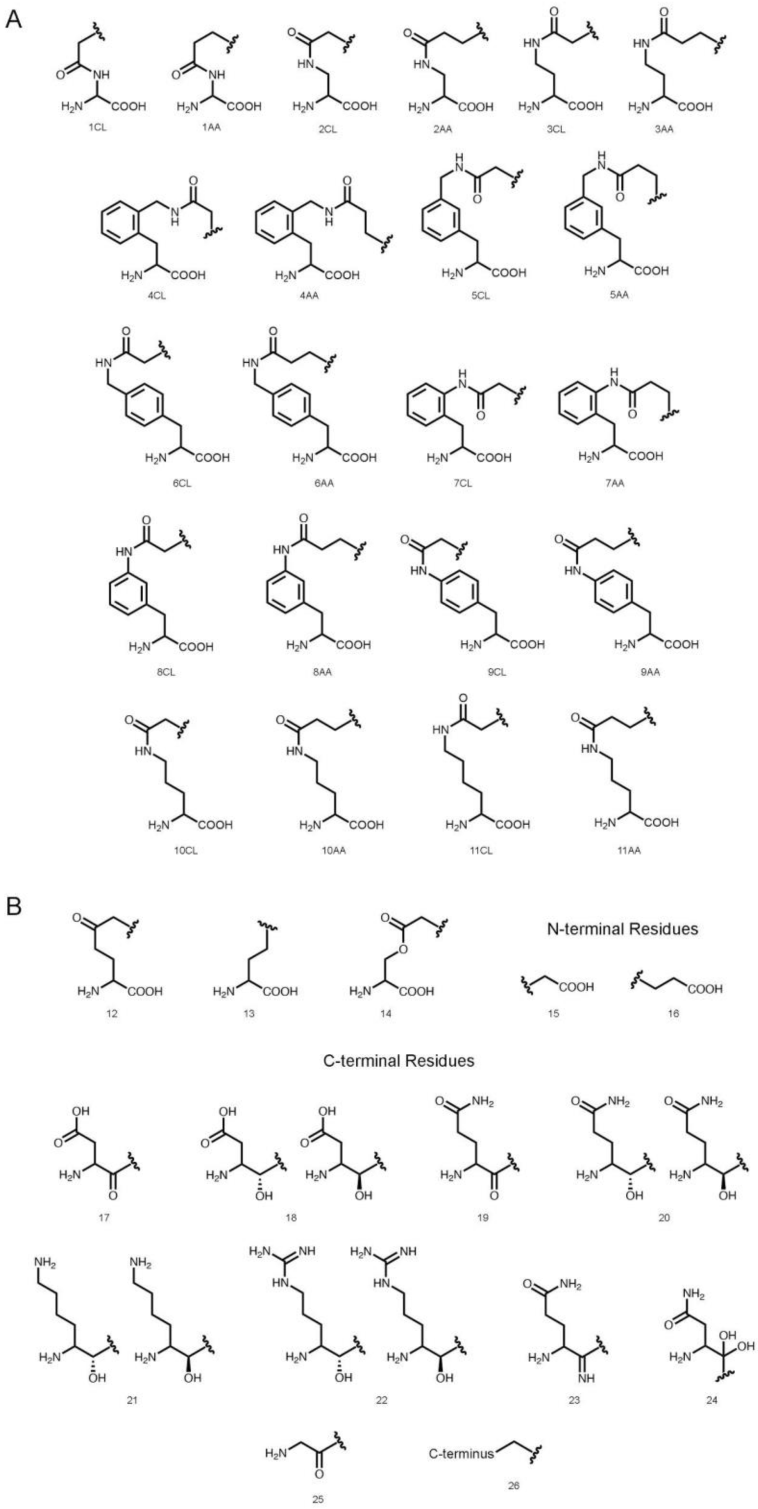


**Supplementary Figure 1: Electrophilic residues parametrized for use in covalent docking, shown in their adduct form. A.** 22 acrylamide- (AA) and chloroacetamide-based (CL) amino acids implemented for use in our design protocol. **B.** Additional 15 electrophilic residues that were implemented to model complexes from the electrophiles dataset.

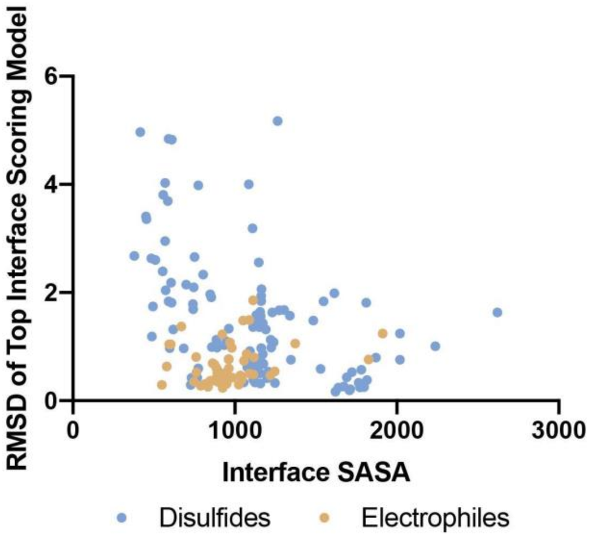


**Supplementary Figure 2: Accuracy as a function of the interface surface area.** The results show a sharp decrease in success rate for complexes with very small interfaces. For example, over the disulfides dataset, the top-scoring model is near-native in only 30% of the structures with SASA < 700, as opposed to 90% of the cases with SASA > 700. Such small interfaces are less common in the electrophiles dataset than in the disulfides dataset (9% and 20%, respectively).

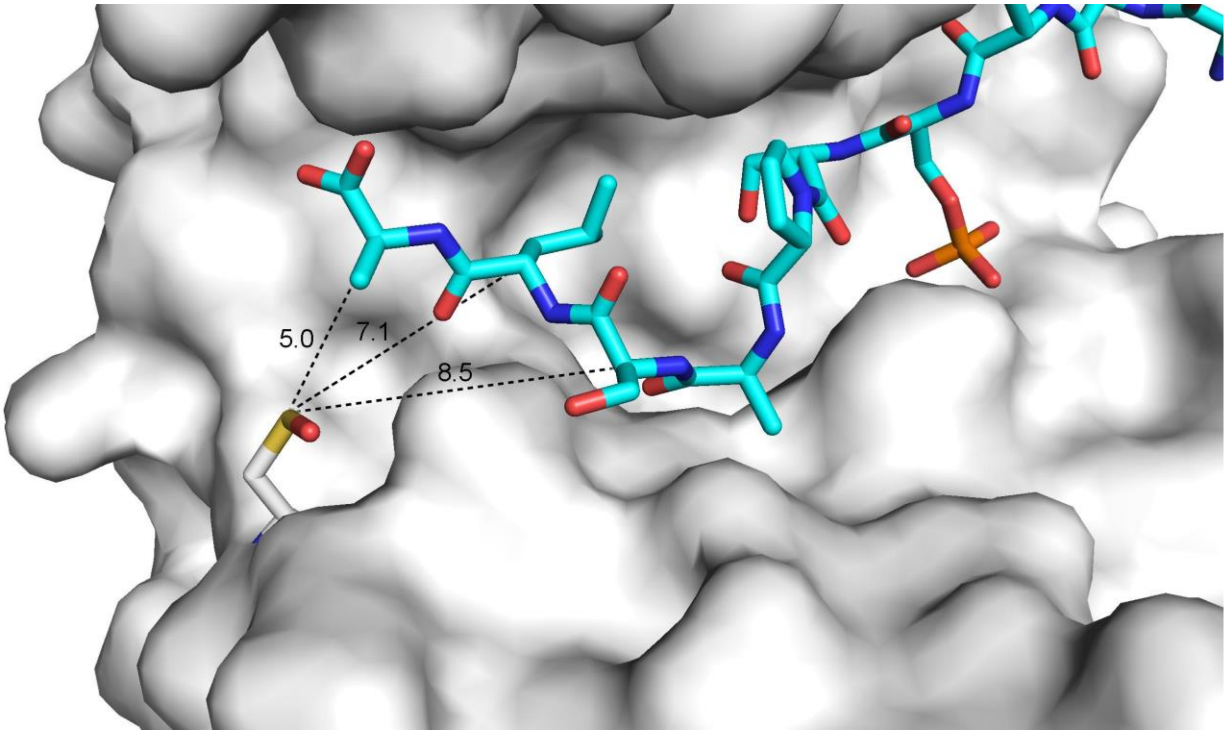


**Supplementary Figure 3: 14-3-3σ non-covalent complex with YAP1 phosphopeptide.** The three C-terminal positions (131-133) were identified as potential sites for electrophile installation. The Cα-Sγ distance is shown in the figure.

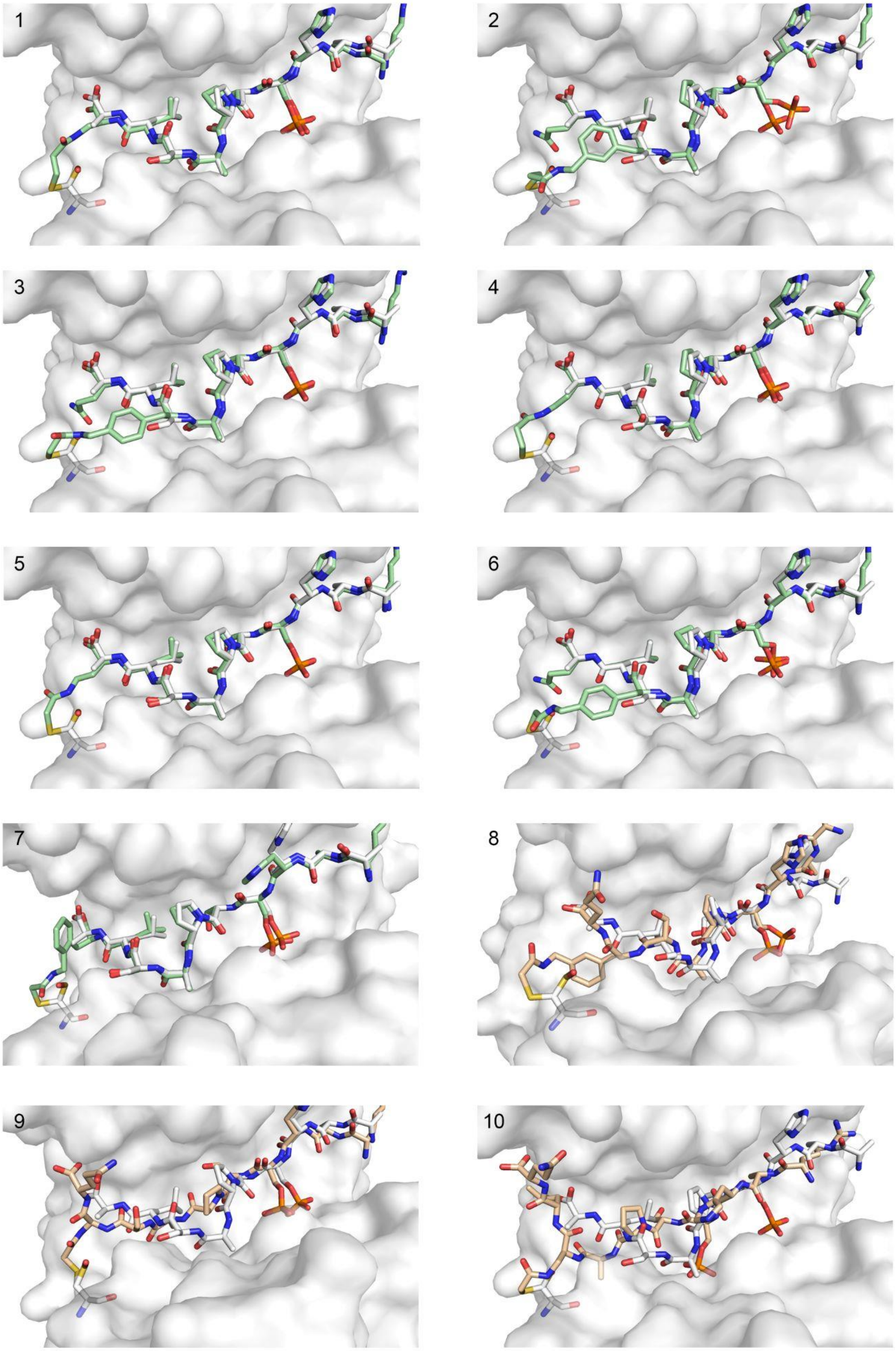
1
2
3
4
5
6
7
8
9
10

**Supplementary Figure 4: 14-3-3σ docking results.** Structural overlay of the docking predictions for peptides 1-10 and the crystal structure of the non-covalent YAP1 phosphopeptide (gray, PDB ID: 3MHR).

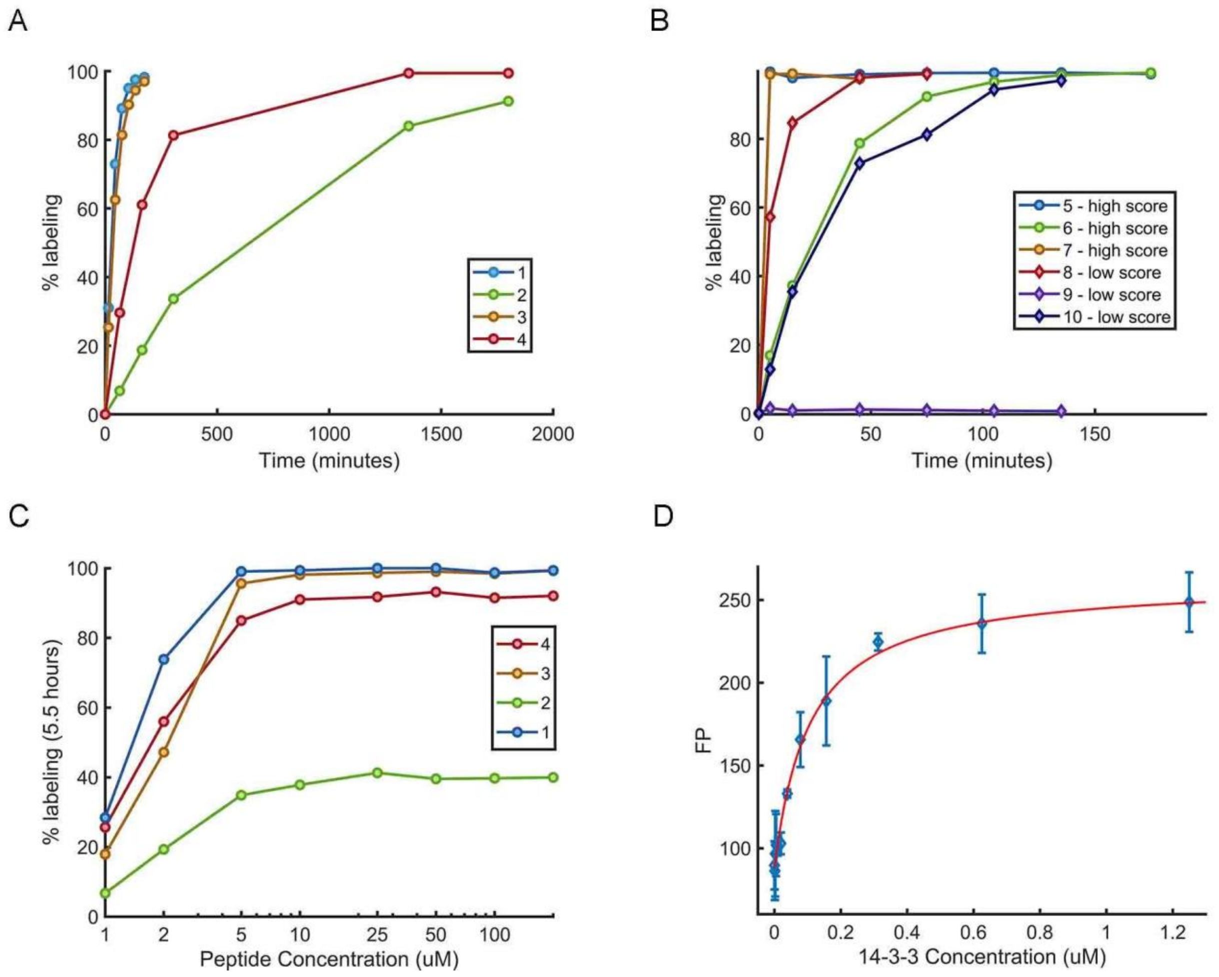


**Supplementary Figure 5:** A) Time course measurement of 14-3-3σ labeling (2 μM) by acrylamide-containing electrophilic peptides (5 μM) at room temperature. B) Time course measurement of 14-3-3σ labeling (2 μM) by chloroacetamide-containing electrophilic peptides (5 μM) at room temperature. C) Dose-response measurement of 14-3-3σ labeling (2 μM) by acrylamide-containing peptides measured at 5.5 hours. D) Fluorescence polarization binding measurement of 10 nM BDP-TMR labeled noncovalent analog of peptide 5

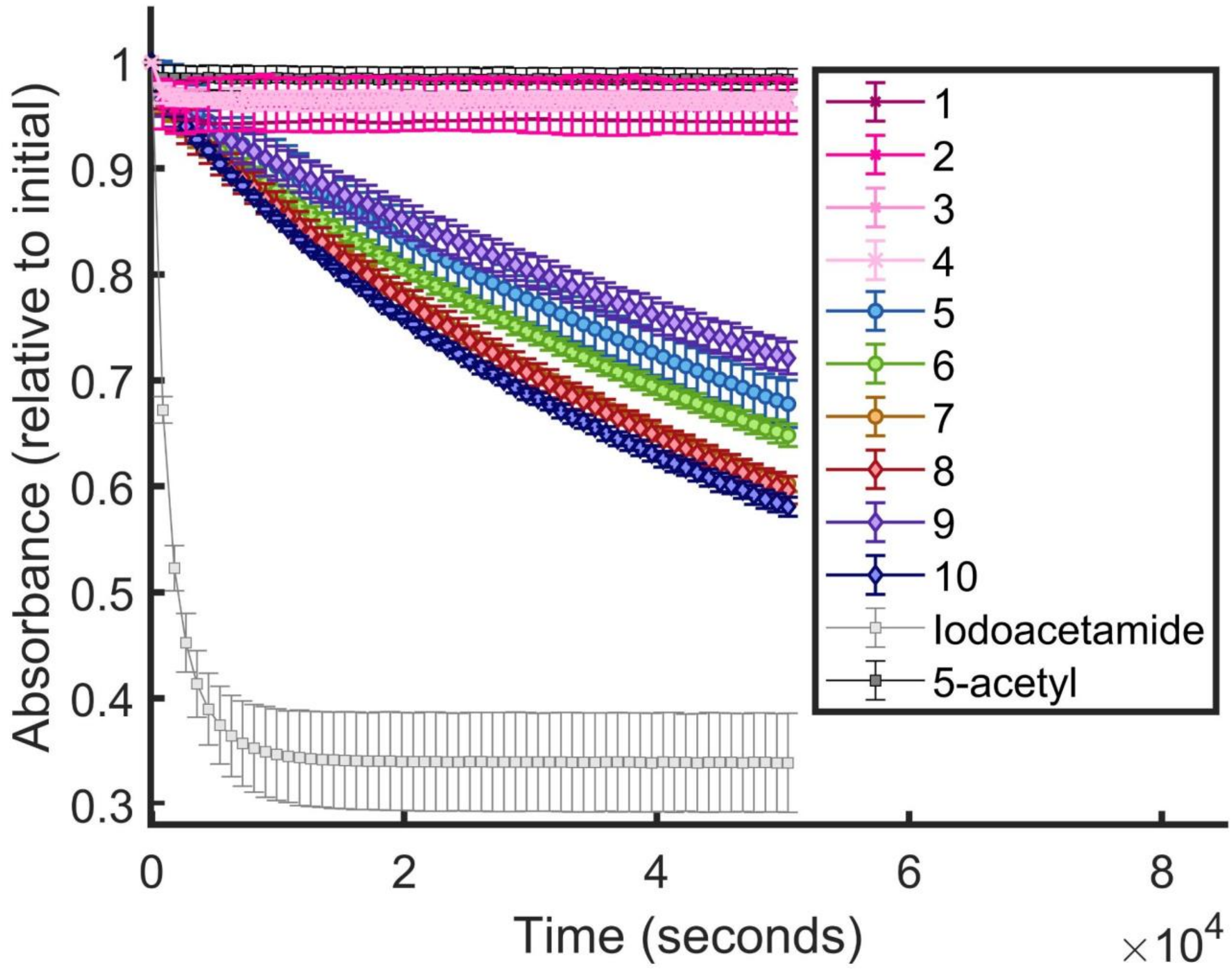


**Supplementary Figure 6:** Thiol reactivity assays of electrophilic peptides 1-10. 200 µM peptides in NaPi 25 mM pH = 7.4, 150 mM NaCl, were reacted with 50 µM DTNB (pre-reduced with TCEP) with monitoring the absorbance at 412 nm every 15 minutes at 37 °C. The acrylamide peptides 1-4, as well as the acetylated control, do not react. The highly reactive iodoacetamide reacts very rapidly, while the chloroacetamide peptides 5-10 display similar reaction rates to one another.

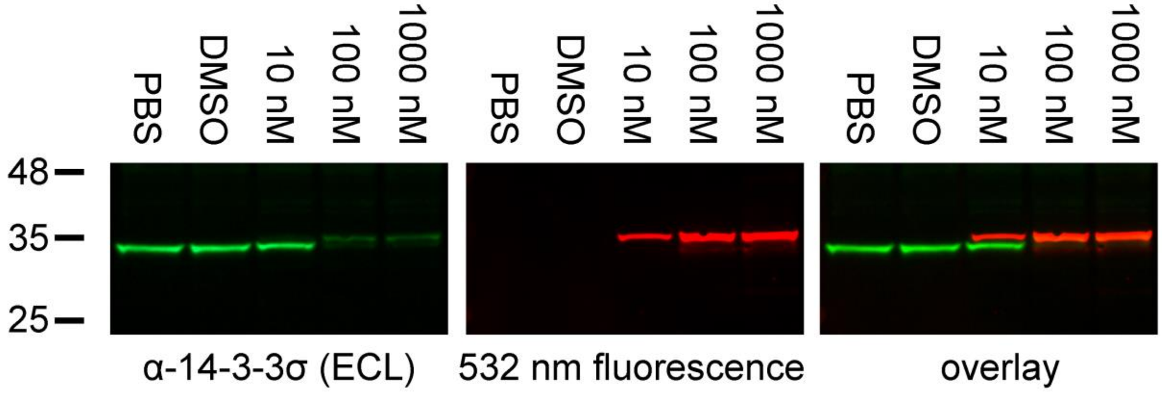


**Supplementary Figure 7:** Validation of target binding using western blot. A431 lysates were incubated with the fluorescent analog of peptide 5, separate on a 4-20% Bis Tris SDS gel and analyzed via western blot. Protein was detected both via anti-14-3-3σ antibody (green, visualized using an HRP-linked secondary antibody through chemiluminescence) and via measurement of fluorescence of the peptide tag (red). Disappearance of the original protein band during peptide binding occurs simultaneously with appearance of the higher mass peptide-protein conjugate, which is detected with significantly decreased intensity by the antibody.

**Table S1: Sequence alignment of the N' terminal region of 14-3-3 proteins**

```
Sigma --M-ERASLIQKAKLAEQAERYEDMAAFMKGAVEKGEELSCEERNLLSVAYKNVVGGQRAAWRVLSSIE
Theta --M-EKTELIQKAKLAEQAERYDDMATCMKAVTEQGAELSNEERNLLSVAYKNVVGGRRSAWRVISSIE
Delta --M-DKNELVQKAKLAEQAERYDDMAACMKSVTEQGAELSNEERNLLSVAYKNVVGARRSSWRVVSSIE
Beta  mtM-DKSELVQKAKLAEQAERYDDMAAAMKAVTEQGHELSNEERNLLSVAYKNVVGARRSSWRVISSIE
Gamma --MvDREQLVQKARLAEQAERYDDMAAAMKNVTELNEPLSNEERNLLSVAYKNVVGARRSSWRVISSIE
Eta   --MgDREQLLQRARLAEQAERYDDMASAMKAVTELNEPLSNEDRNLLSVAYKNVVGARRSSWRVISSIE
Epsil --MdDREDLVYQAKLAEQAERYDEMVESMKKVAGMDVELTVEERNLLSVAYKNVIGARRASWRIISSIE
```

**Table S2.** Data collection and refinement statistics (molecular replacement) for 14-3-3σ bound to peptide 6 (PDB: 7O07)

| | 14-3-3σ in complex with peptide 6 |
|---|---|
| *Data collection* | |
| Space group | C 2 2 21 |
| Cell dimensions | |
| a, b, c (Å) | 82.6, 112.6, 63.2 |
| α, β, γ (°) | 90, 90, 90 |
| Resolution (Å) | 66.59 (1.20) (1.22 – 1.20) |
| $I$ / σ($I$) | 11.8 (1.9) |
| Completeness (%) | 100.0 (100.0) |
| Redundancy | 12.2 (11.4) |
| $CC_{1/2}$ | 0.998 (0.785) |
| *Refinement* | |
| No. reflections | 91999 |
| $R_{work}/R_{free}$ | 0.184/0.1959 |
| No. atoms | |
| Protein | 2064 |
| Ligand/ion | 21 |
| Water | 316 |
| *B*-factors | |
| Protein | 14.81 |
| Ligand/ion | 16.70 |
| Water | 27.33 |
| R.m.s. deviations | |
| Bond lengths (Å) | 0.004 |
| Bond angles (°) | 0.72 |
